# Functional Network Community Detection Can Disaggregate and Filter Multiple Underlying Pathways in Enrichment Analyses

**DOI:** 10.1101/166207

**Authors:** Lia X. Harrington, Gregory P. Way, Jennifer A. Doherty, Casey S. Greene

## Abstract

Differential expression experiments or other analyses often end in a list of genes. Pathway enrichment analysis is one method to discern important biological signals and patterns from noisy expression data. However, pathway enrichment analysis may perform suboptimally in situations where there are multiple implicated pathways – such as in the case of genes that define subtypes of complex diseases. Our simulation study shows that in this setting, standard overrepresentation analysis identifies many false positive pathways along with the true positives. These false positives hamper investigators’ attempts to glean biological insights from enrichment analysis. We develop and evaluate an approach that combines community detection over functional networks with pathway enrichment to reduce false positives. Our simulation study demonstrates that a large reduction in false positives can be obtained with a small decrease in power. Though we hypothesized that multiple communities might underlie previously described subtypes of high-grade serous ovarian cancer and applied this approach, our results do not support this hypothesis. In summary, applying community detection before enrichment analysis may ease interpretation for complex gene sets that represent multiple distinct pathways.

## 1. Introduction

Researchers’ experiments that include high-throughput data generation often lead to a set of genes. These genes may be genes that are over- or under-expressed in a disease subtype, are upregulated in response to a drug, or contain variants associated with a disease. After potentially interesting genes are identified, the next challenge is to interpret the biological processes or pathways that underlie the set. Overrepresentation-based methods are commonly used to identify pathways that have more members in the identified set than would be expected by chance^1^. Typically, pathways or similar groups of genes are obtained from structured vocabularies outlined in curated ontologies such as KEGG, PID, GO, or Reactome^2–5^. Recently, computational researchers have sought to improve the power of such analyses by considering network interactions among pathway members^6,7^. We sought to evaluate overrepresentation analysis in a different setting: one where multiple pathways underlie a set of associated genes. In this situation, applying standard overrepresentation analysis to gene sets constructed by randomly selecting members of multiple pathways identifies many false positive pathways. We hypothesized that reducing the noise of the gene list input via community detection might decrease the number of false positive pathways.

Functional networks are a type of network where genes are connected if they have a high probability of working together in the same pathway or process^8-11^. To address the challenge posed by multi-pathway gene sets, we developed an approach that incorporates information from functional networks to first partition gene sets into subsets, or communities, which are then analyzed for overrepresented pathways. To accomplish this, enrichment analysis is applied to each extracted community resulting from community detection preprocessing^12,13^ of the original gene set. Community detection has been applied to financial data, social media, and biological data^12,14^.To our knowledge, this is its first application to disambiguate the pathways associated with complex gene sets. We evaluate four community detection methods in this context: Fastgreedy, Walktrap, Multilevel, and Infomap. These algorithms all aim to identify groups/communities within a network:

- Fastgreedy – This algorithm starts from a completely unclustered set of nodes and iteratively adds communities such that the modularity (score maximizing within edges and minimizing between edges) is maximized until no additional improvement can be made^15^.
- Walktrap – This algorithm performs random walks using a specified step size. Where densely connected areas occur, the random walk becomes “trapped” in local regions that then define communities^16^.
- Multilevel – This algorithm is similar to fastgreedy, but it merges communities to optimize modularity based upon only the neighboring communities as opposed to all communities^17^. The algorithm terminates when only a single node is left, or when the improvement in modularity cannot result from the simple merge of two neighboring communities.
- Infomap – This algorithm uses the probability flow of information in random walks, which occurs more readily in groups of heavily connected nodes. Thus, information about network structure can be compressed in maps of modules (nodes where information travels quickly)^18^.

Outside of the multi-pathway gene set challenge, there are a number of R packages that implement algorithms for network interpretation of experimental results including WGCNA^19^, EnrichNet^20^, pathDIP^21^, and CePa^22,23^. In this work, community detection algorithms are used to partition multi-pathway gene sets before overrepresentation analysis. By detecting these gene communities, we aim to provide cleaner inputs for overrepresentation analyses in the case of multiple underlying pathways – thereby reducing the number of identified false positives. In contrast with other methods that use network information as priors or as post-analysis visualization aides, we group genes before enrichment analysis. While we use the Integrative Multi-species Prediction (IMP) networks, our approach can be applied to a gene set from any source^11,24^. For example, a user may wish to use tissue-specific networks from the GIANT webserver^9^ if tissue specificity is important. Finally, our approach makes no assumptions about the covariance structure of the networks^25^ and is thus potentially more useful in real world applications where certain assumptions may not apply.

In summary, we propose an alternative gene enrichment approach for cases when multiple pathways are suspected to be implicated in a gene list. In this approach, candidate genes are overlaid onto a functional network and separated into communities of related genes via community detection. Communities are then subjected to an overrepresentation analysis independently and multiple testing corrections are applied. We compare four community detection approaches in simulated experiments and then apply the approach to identifying enriched pathways across high grade serous ovarian cancer (HGSC) subtypes.

## 2. Methods

We conducted an experiment that contained a control and an experimental arm. The control arm was an overrepresentation analysis without community detection, and the experimental arm was an overrepresentation analysis with various community detection methods applied as a preprocessing step.

### 2.1. General Approach

From the KEGG ontology, *m* randomly chosen pathways were selected to form a list of candidate genes. To help evaluate the impact of incomplete pathway discovery, only *p* percent of the genes in each pathway were randomly selected for inclusion in the final gene list. Finally, *a* percent of additional random genes selected without replacement from the ontology were added to the gene list to create noise. As to only consider genes that influence pathway analysis, genes that were not in both IMP and KEGG were excluded for a resulting set of 5195 genes. This procedure was performed for both control and experimental arms so that differences in results could be attributed to community detection preprocessing.

We performed one hundred iterations for each parameter level combination of number of pathways (*m* = 2-8), percentage of genes included from each pathway (*p* = 30%, 47.5%, 65%, 82.5%, and 100%), and percentage additional random genes from IMP (*a* = 10%, 32.5%, 55%, 77.5%, and 100%) for a total of 105,000 individual runs. Over the 100 iterations of the specific parameter combination, we measured the number of seeded pathways correctly detected (true positives), incorrectly detected (false positives), correctly missed (true negatives), and incorrectly missed (false negatives). The false positive proportion, false negative proportion, precision, recall, and F1 score were calculated for each parameter combination over the 100 iterations. The F1 score is the weighted average of precision and recall where precision is the number of true positives divided by all positives and recall is the number of true positives divided by the sum of true positives and false negatives.

### 2.2. Control Arm

The control arm followed the steps outlined in General Approach.

#### 2.2.1. Control All (CtrAll)

For this method, we determined true positives, false positives, true negatives, and false negatives using all significantly enriched pathways and complete gene lists of seeded pathways. For example, if a gene list was seeded with three pathways and the enrichment analysis identified ten pathways (including correctly identifying the original three), then all ten pathways would be counted as positives with the seven unseeded pathways considered false positive.

#### 2.2.2. Control M (CtrM)

For this method, true positives, false positives, true negatives, and false negatives were determined using only the top *m* significant pathways where *m* is the number of seeded pathways. For example, if three pathways were seeded and there were ten significant pathways, then only the top three pathways in the significant enrichment results would be considered. Thus, if all three seeded pathways were in the top three significant results, the true positive would be three and false positive would be zero. If, however, only two of the three seeded pathways were in the top three significantly enriched pathways, then true positive would be two and false positive would be one. CtrM provides provides an upper bound on possible performance as it is unrealistic in practice for investigators to know *a priori* the correct number of pathways.

### 2.3 Experimental Arm

For the experimental arm, the subgraph associated with each gene list described in the General Approach was extracted from IMP and subjected to community detection to provide community-level gene sets before the overrepresentation analysis. Fastgreedy, Walktrap, Infomap, and Multilevel community detection algorithms were applied in the community detection step. The communities of genes detected by the algorithm were then used as separate candidate gene lists for overrepresentation analysis. True positive, false positive, true negative, and false negative were calculated for all pathways that remained statistically significant after Bonferroni multiple testing correction at α = .05 was applied. This correction was applied for each community if multiple were found.

All simulation analyses were performed using Python 2.7.6 with the iGraph package (version 0.71). Figures were produced using ggplot in R 3.3.1. Open source software to reproduce the results of this paper is provided at https://github.com/greenelab/GEA_Community_Detection. Figure 1 provides an overview of both the control and experimental arms.

**Fig. 1.**
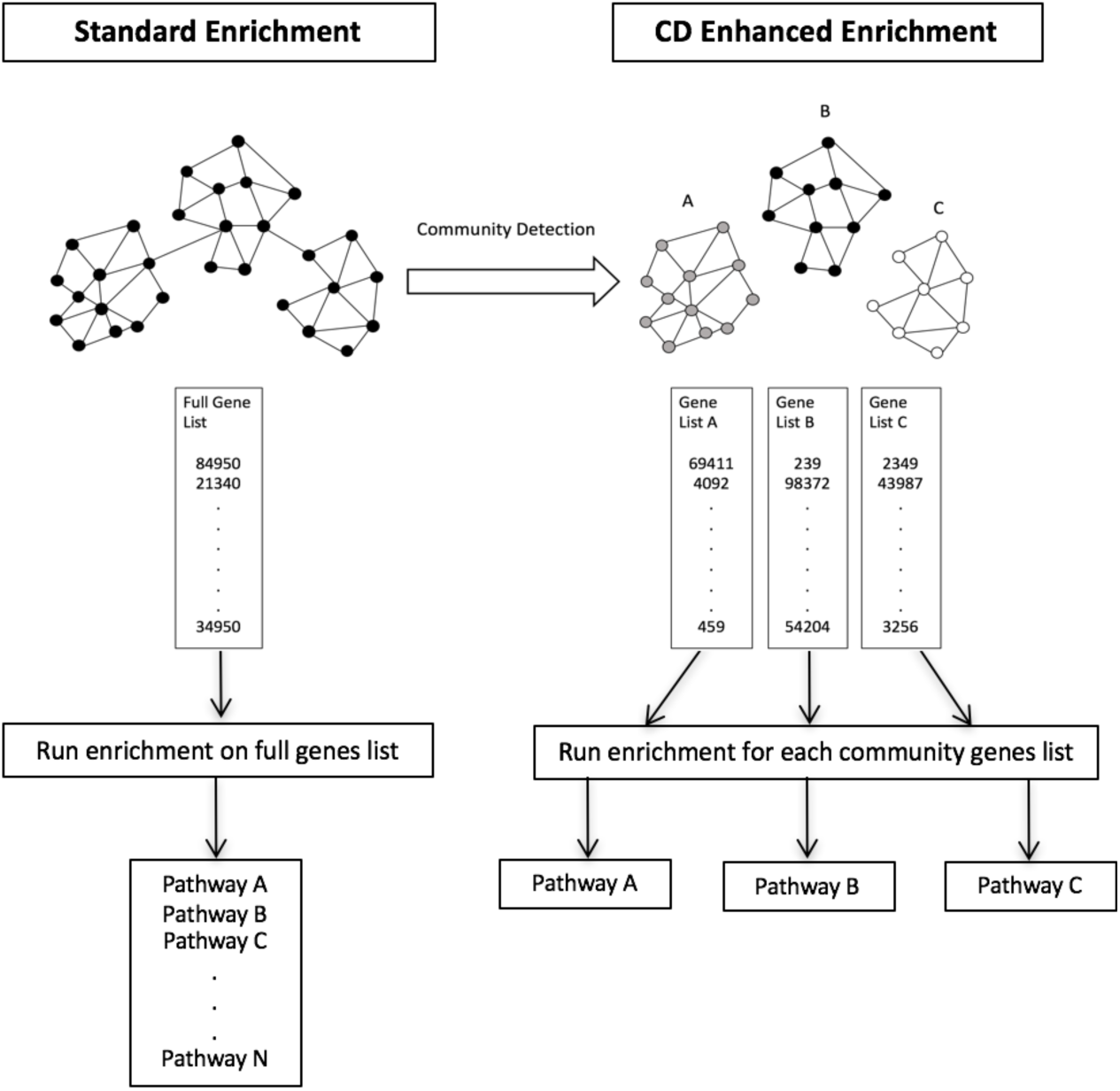
In standard enrichment analysis, the full gene list is subjected to enrichment analysis and all significantly enriched pathways are returned. In the proposed experimental community detection enhanced method, the full gene list is first subjected to community detection to parse the gene list into sub-gene lists. Enrichment analysis is then performed for each gene list associated with each “discovered” community. Only the most significant pathway is returned for each community.

### HGSC Application

Based on the results of the simulation study, we applied the top performing community detection algorithms to lists of genes characterizing high-grade serous ovarian cancer (HGSC) subtypes. The gene lists were previously identified by a one cluster versus all differential expression analysis^26^ of cluster specific genes in common to four HGSC datasets^27–30^. While previous reports have described four HGSC subtypes, the multi-population study suggested that the number was three or fewer^26^. Given these conflicting results, we applied community detection to HGSC subtype-specific gene lists previously derived from results classifying 2, 3, and 4 subtypes^26^. Because this is an analysis of cancer genomics data, we used cancer pathways from the Pathway Interaction Database (PID)^5^.

## 3. Results and Discussion

### 3.1. Simulation Study

In general, community detection methods reduced the number of false positive associations in the multi-pathway setting. When seeding a gene list with four random pathways, all community detection methods had higher F1 scores than the standard enrichment analysis, CtrAll (Figure 2). In cases where pathways were incompletely seeded, the community detection methods often outperformed CtrM, which only considers the top *m* pathways as statistically significant (Figure 2). These findings are consistent when using the top 2-8 pathways (pathway numbers 2, 3, 5, 6, 7, and 8 are Supplementary Figures S1-6). Performance was robust to the number of genes taken from each seeded pathway over a broad range of values, and the relative performance of methods was largely unaffected by the proportion of genes sampled from the seeded pathways (i.e, 30% or all 100%) to make the gene lists. Thus, our approach may be more useful than standard enrichment techniques in situations where one is presented with a long, heterogeneous, and incomplete gene list and one wishes to find a set of robust pathways for further investigation. The Walktrap and Multilevel methods demonstrated the most success in this context as they resulted in high F1 scores and relatively low false negative and false positive proportions. Compared to other community detection methods, Fastgreedy appeared to have a broader range of performance values, with higher variability and increased outliers. The performance of community detection algorithms may be network-specific; users may wish to apply our open source code to perform a new simulation study if different networks are selected.

**Fig. 2.**
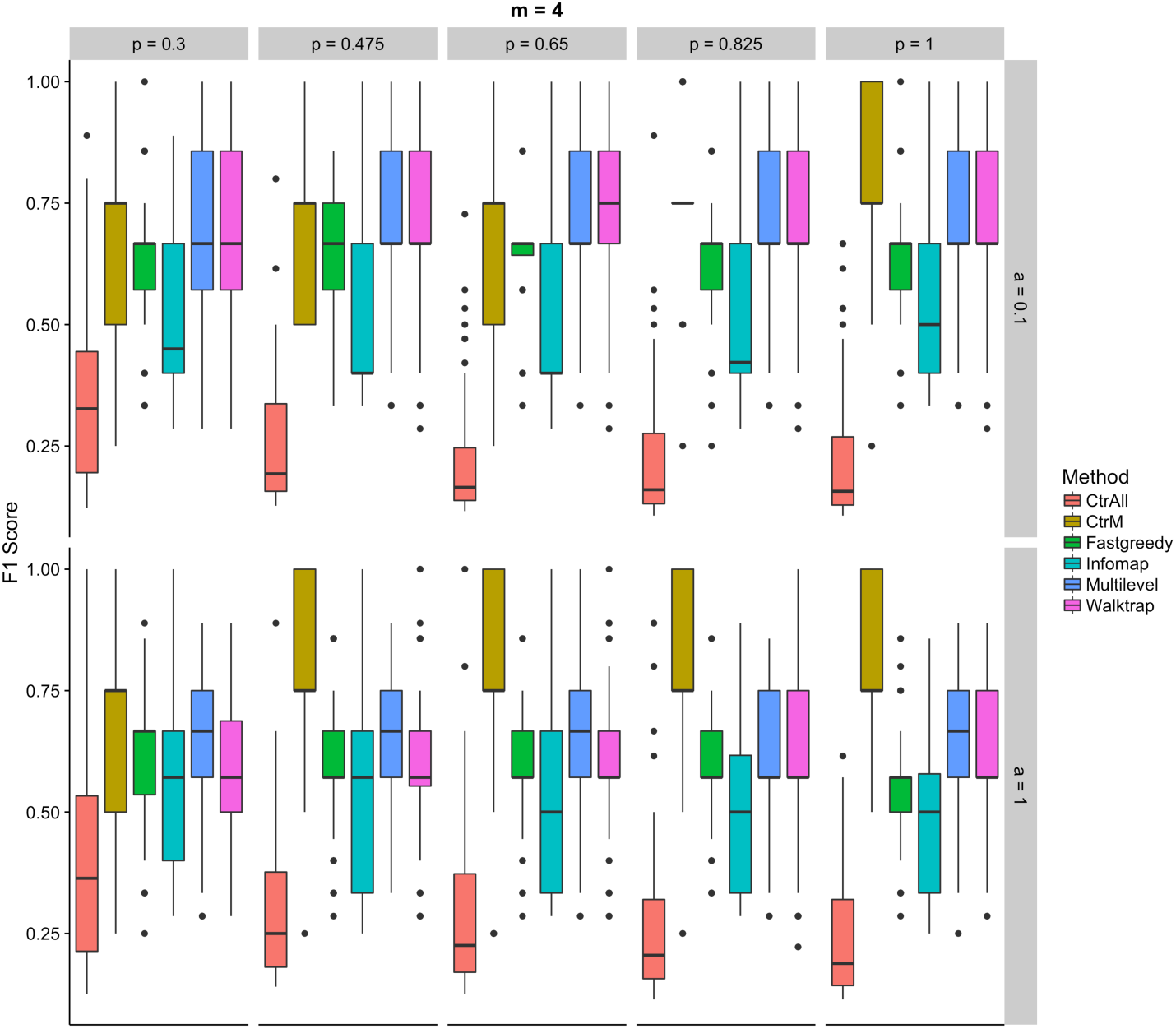
F1 scores for the controls (using all (CtrAll), or only the top 4 (CtrM), statistically significant pathways) and the community detection methods: Fastgreedy, Infomap, Multilevel, and Walktrap for various percentages of genes in each pathway (top axis) and percentages of additional genes (right side axis) for simulations using 4 random pathways. The percentage of genes indicates the percentage of random genes selected from each pathway. The percentage of additional genes indicates how many unrelated genes are randomly added to the analysis to represent increasing amounts of noise. Each comparison includes 100 iterations.

The combination of community detection and enrichment was designed to filter false positives in the multi-pathway setting. When we evaluated the proportion of false positives, we observed that the F1 score improvements were driven by successful filtration. In particular, all community detection methods outperformed standard enrichment analyses for false positive proportions (Figure 3). As expected, when the number of seeded pathways increased, the proportions of false positives steadily increased for control runs that included all statistically significant pathways. The standard enrichment analysis approach was well suited to identifying a single pathway. The more pathways that were present in a single genelist, the worse standard enrichment-based methods performed.

**Fig. 3.**
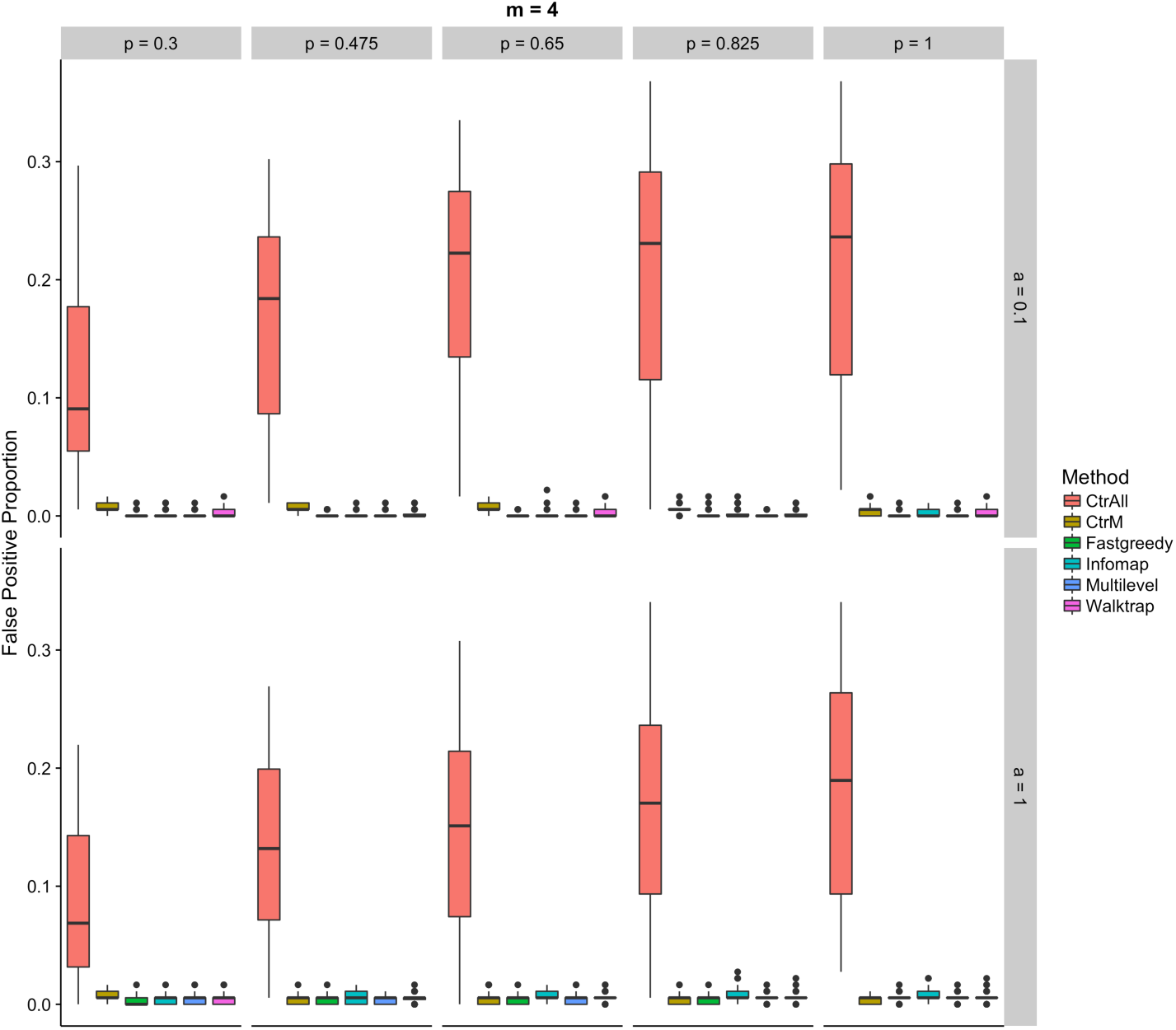
Proportions of false positives for the controls (using all (CtrAll), or only the top 4 (CtrM), statistically significant pathways) and the community detection methods: Fastgreedy, Infomap, Walktrap, and Multilevel for various percentages of genes in each pathway (top axis) and percentage of additional genes (right side axis) for simulations using 4 random pathways.

All community detection methods other than CtrAll usually miss some portion of the true positives using 4 seeded pathways (Figure 4). In general, Walktrap, Infomap, and Multilevel tend to have greater variability in the number of pathways missed compared to CtrAll and Fastgreedy. It is not surprising that the community detection and CtrM methods have higher proportions of false negatives than CtrAll since they were designed to reduce false positives. Thus, a traditional enrichment approach may be more appropriate in sitatuions where false negatives are more of a concern, such as when investigating a relatively small gene list or conducting an exploratory analysis.

**Fig. 4.**
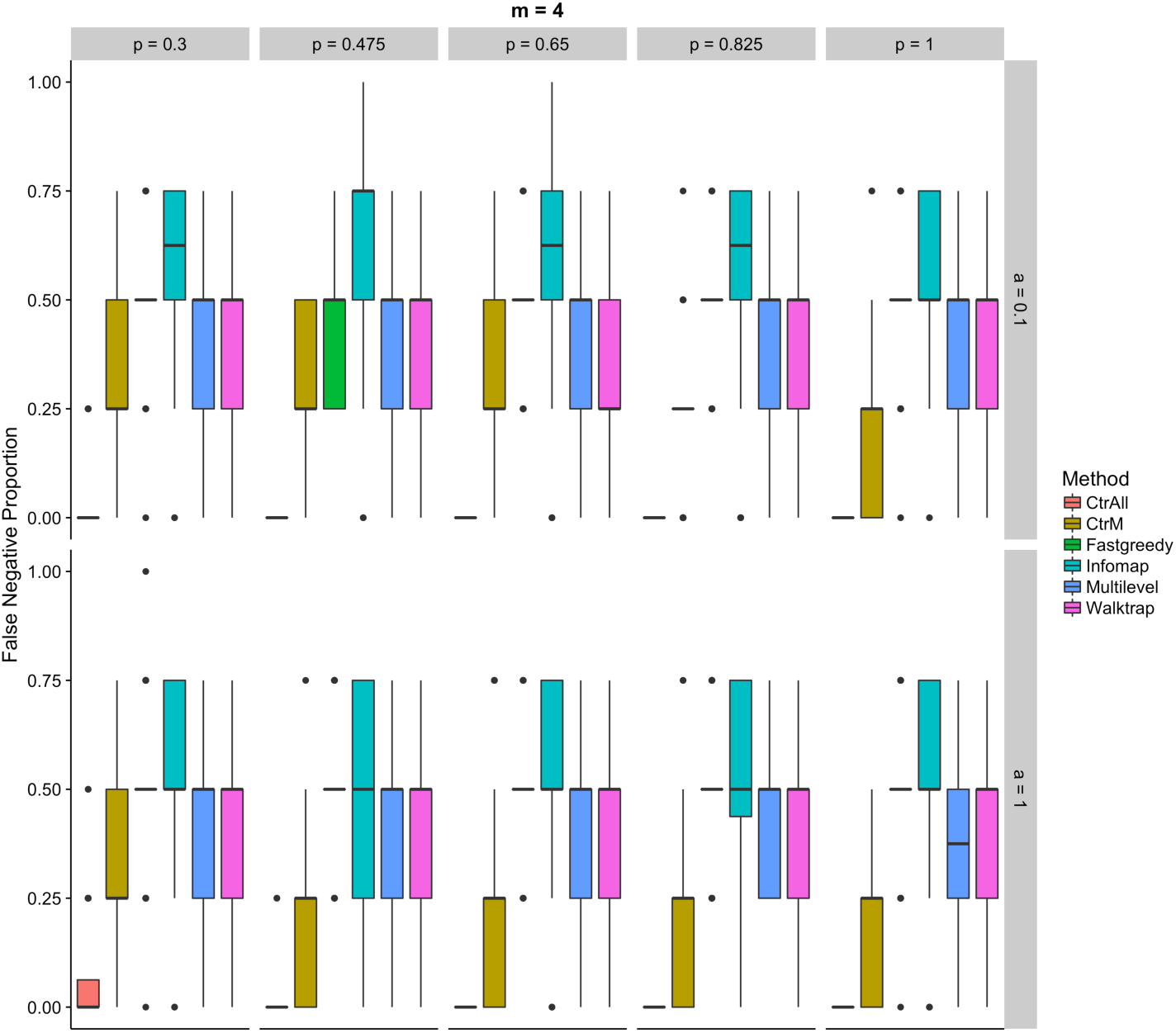
Proportions of false negatives in the controls (using all (CtrAll), or only the top 4 (CtrM), statistically significant pathways) and the community detection methods: Fastgreedy, Infomap, Walktrap, and Multilevel for various percentage of genes in each pathway (top axis) and percentage of additional genes (right side axis) for simulations using 4 random pathways.

### 3.2. HGSC Results

To examine the biological applicability of community detection, we independently applied the community detection approach to previously defined, HGSC subtype-specific gene lists for when 2, 3, and 4 subtypes are assigned. We previously derived these gene lists from a differential expression analysis across HGSC subtypes that were concordant across different populations^26^. We selected only the top performing algorithms from our simulation study, Walktrap and Multilevel. Applying these methods to PID pathways, we found that most clusters mapped to either Beta1 integrin cell surface interactions or IL12-mediated signaling events (Table 1). Community detection methods was able to separate upregulated and downregulated genes coming from the same pathway into different communities (Table 1)

**Table 1.**
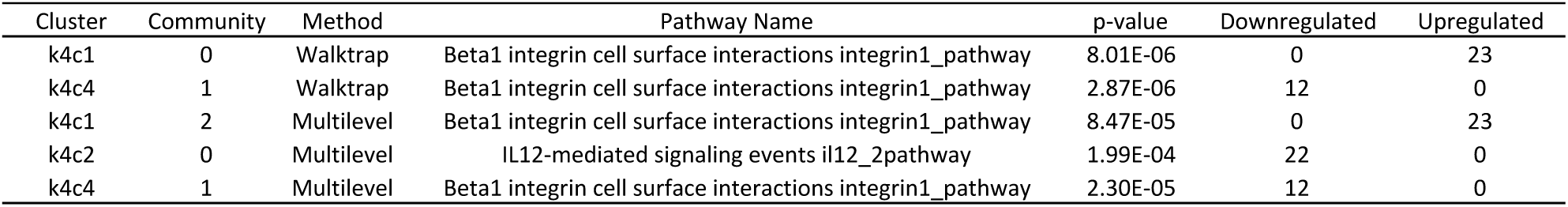
The statistically significantly enriched pathways found by Walktrap and Multilevel community detection methods and the number of genes in each pathway that are either upregulated (more highly expressed) or downregulated (less expressed) in HGSC^26^. We identified statistically significant pathways in communities defined by only k = 4 in cluster 1 (k4c1), cluster 2 (k4c2), and cluster 4 (k4c4). The id number of the enriched community is also provided. Clusters 1, 2, 3 and 4 correspond to mesenchymal, proliferative, immunoreactive, and differentiated subtypes as previously defined by TCGA^27^

While many pathways were implicated in the original pathway analysis (see Supplementary Table S6 of Way et al. 2016^26^), our community detection approach only implicated two distinct pathways consistently, for 2-4 subtypes. This did not support our hypothesis that HGSC subtypes are driven by differences across multiple pathways that are captured in differentially expressed gene lists. HGSC subtypes are known to be primarily characterized by a mesenchymal gene signature and immunoreactivity. Our analysis suggested that up- and down-regulation of beta 1 integrin signaling, and down-regulation of IL12 signaling, primarily define the subtype-specific signatures. However, the lack of PID pathway enrchiment in the presence of community structure may indicate novel biological pathways driving subtype separation. Beta 1 integrin signaling is a well characterized indicator of metastasis^31^ and its high expression is associated with poor survival in ovarian cancer patients^32^. IL12 is an important immune system process with many coordinated functions^33^. Importantly, administration of intraperitoneal IL12 is being explored as a therapeutic agent in ovarian cancer^34^. The community detection approach pointed to specific HGSC subtypes that were aligned with this characterization, but did not identify multiple pathways for any specific subtype. We often observed that pathways that were highly expressed for one subtype would be underexpressed for another, which was consistent with a model that HGSC subtypes exist along a continuum of underlying pathway or cell type content. These results are also generally consistent with those found previously^27, 28,35, 36^.

## 4. Conclusion

In summary, we developed an alternative enrichment method that uses community detection to group genes based on network connectivity prior to enrichment analyses. This approach is designed for situations where a researcher hypothesizes that multiple pathways contribute to a gene set. It trades an increase in false negatives for a dramatic reduction in false positives. The standard enrichment approach may be more appropriate in exploratory stages of research when high power is more desired than false positive control. Applying this method to gene sets that characterize HGSC subtypes did not reveal multiple pathways underlying any of the previously described subtypes. These results are consistent with a model where factors other than the activity of multiple pathways are responsible for the difficult to discern HGSC subtypes.

## 5. Acknowledgments

We thank James Rudd for several thoughtful conversations about the developed approach. This work was funded in part by grants from the Gordon and Betty Moore Foundation (GBMF 4552) to CSG and from the National Institutes of Health (R01 CA200854) to JAD and CSG.

## 6. Supplementary Material

Supplementary figures can be found at http://doi.org/10.5281/zenodo.830568^37^.

